# *Clostridium butyricum* and *Clostridium tyrobutyricum*: angle or devil for Necrotizing enterocolitis?

**DOI:** 10.1101/2023.04.10.536220

**Authors:** Ruizhi Tao, Gangfan Zong, Yehua Pan, Hongxing Li, Peng Cheng, Rui Deng, Wenxing Chen, Aiyun Wang, Shishan Xia, Weibing Tang, Yin Lu, Zhonghong Wei

## Abstract

**Background:** Necrotizing enterocolitis (NEC), high incidence and case-fatality rate among premature neonates, is a frustrating gastrointestinal disease which hard to eradicate currently for its unclear pathogenesis and mechanisms. What has been conformed is that the gut microbes dysbiosis happens before the occurrence of NEC, providing robust evidence for the usage of probiotic therapy. Hence, we mainly concentrated on two probiotics: Clostridium butyricum and Clostridium tyrobutyricum especially after the breakthrough in discovering that several clostridia species have associations with NEC.

**Result:** To verify whether these two clostridia are pathogenic or probiotic, we compared the phenotypic traits of NEC mice treated with two clostridia. Our results proof that treatment with C. tyrobutyricum recovers the intestinal barrier integrity and alleviates inflammatory immune response of NEC, while treatment with C. butyricum aggrevates the intestinal barrier damage and promotes immune disorder including the number of macrophages, monocytes and neutrophils in Intestinal lamina propria. Further analysis of gut micrbiome implies that the positive effect of C. tyrobutyricum treatment is in association with the increase of Akkermansia muciniphila. Meanwhile, C. butyricum treatment decreases the level of A. muciniphila, which accounts for the negative effect to NEC.

**Conclusion:** This study sheds light on that treatment with C. tyrobutyricum but not C. butyricum is entitled to protect against NEC development potentially. The mechanisms behind the opposite effect on NEC may result in different modulation on the level of A. muciniphila, which is deeply associated with intestinal homoeostasis. Briefly, through improving the abundance of A. muciniphila to alleviate intestinal inflammation and enhance intestinal barrier integrity, C. tyrobutyricum supplement may become a promising therapy for NEC.

## Background

Necrotizing enterocolitis has become the most severe and common acquired gastrointestinal diseases in prematurity [1] and up to 12% premature infants with an extremely low birth weight (VLBW<1500g) are more likely to catch NEC [2]. Not only antenatal risk factors such as intrauterine inflammation, infection and preeclampsia [3], but also postnatal risk factors including prematurity, sepsis, formula feeding and gut microbes dysbiosis account for the development of NEC[4]. Limited to inadequate acknowledgment about the complex and multifactorial pathogenesis and mechanisms, the mortality of NEC neonates remains fluctuating between 20%∼30% even after surgery[5].

Nevertheless, a mass of literatures available declare that the imbalance of gut microbiota plays a critical role in NEC, which precedes the occurrence of the disease[6]. Meanwhile, early bacterial colonization is vital for the formation of intestinal barrier integrity and systemic immune function in infants [7]. Aberrant chemical and physical barriers of intestine leads to bacteria invasion, thus perforation, necrosis and systemic inflammatory reaction occurring [8]. Therefore, gut dysbiosis is a core risk factor to be reckoned with during the development of NEC and probiotic therapy is emerging as the times require[9, 10].

Metagenomic analysis indicates an increased relative abundance of phylum Proteobacteria and a decreased relative abundance of phylum Firmicutes and Bacteroidetes in NEC infants [11]. What is noteworthy now is that whether genus Clostridium is positive or negative in association with NEC remains obscure, rendering a new puzzle for NEC therapeutic strategies. Some investigators reported that Clostridium butyricum [12], Clostridium perfringens [13] and Clostridium neonatale [14] are positive strain-specific associated with NEC. However, one literature pointed out a decreased Clostridia abundance with the development of NEC [15].

Hence, we focus on two probiotics of clostridium: Clostridium butyricum and Clostridium tyrobutyricum, which are both butyric acid-producing bacteria and have been applied to treat gastro-enteritis clinically or experimentally [16–18]. To figure out whether they are pathogenic or probiotic for NEC, we compared phenotypic traits, intestinal barrier integrity and inflammatory immune response of NEC in a mouse model treated with the two Clostridia. And results show that Clostridium tyrobutyricum alleviates the symptoms of NEC while Clostridium butyricum promotes the development of NEC. Furthermore, we found that the abundance of Akkermansia muciniphila was enhanced by Clostridium tyrobutyricum treatment and weakened by Clostridium butyricum treatment through 16S rDNA analysis, which implied the interspecific competition and colonization Resistance between them. Together, our data advance knowledge on the screening of potential probiotics for clinicians to treat NEC with multiple strategies and highlight the significance of early bacterial colonization for infants and the potential interspecific competition mechanisms among these bacteria.

## Materials and methods

### Human samples collection

Human samples including colons and feces in the sterile surgery room were collected from the Jiangning affiliated Hospital of Nanjing Medical University and Nanjing Children’s Hospital respectively. Colon samples were transferred by germ free 50ml Eppendorf tubes which contained with sterile RPMI Medium 1640 and the feces samples were transferred by germ free collection tubes. All transportations were under dry ice condition for further study. The information of all human samples was provided in Table S1.

### Bacterial Strains and Culture Conditions

Clostridium butyricum (ATCC 19398) and Clostridium tyrobutyricum (ATCC 25755) and were purchased from American Type Culture Collection (ATCC, Manassas, USA). Akkermansia muciniphila (DSM 22959) was purchased from Deutsche Sammlung von Mikroorganismen und Zellkulturen (DSMZ, Braunschweig, Germany). Both Clostridia were cultured in Modified Reinforced Clostridium Medium (RCM) broth (M1285-01, ELITE-MEDIA) at 37°C and Akkermansia muciniphila (DSM 22959) was cultured in Brain Heart Infusion (BHI) broth (Hopebio, HB8297-5) with0.1% mucin (SIGMA, M2378-100G) in aerobic conditions.

### Mouse Model of Necrotizing Enterocolitis with or without Treatment

Pregnant C57/BL6 mice were purchased from Jiangsu Jicuiyaokang Biotechnology Co., LTD. When newborn mice were 1 week old, mice were randomly divided into different groups. Control mice were fed with breast milk without separated from their dams. NEC mice were co-housed in a neonatal incubator at 32-35°C and 60-70% humidity. They were hand-fed with formula milk (40μl/g), consisted of 15g Similac Advance infant formula (Abbott Nutrition) and 10g Esbilac canine milk replacer (PetAg) in 75ml water with or without bacteria (12.5μl stool slurry/ml or 10^9^ colony-forming units/ml) 6 times a day. Meanwhile, they were stressed with hypoxia (5%O_2_, 95%N_2_) for 10 min, followed by cold stress (placement in a refrigerator at 4 °C) for 5 min later twice a day. Mice were humanely sacrificed after 1 week and the intestines were collected immediately for further experiments. Shortly, Control group: breast-fed only. NEC group: formula-fed, hypoxia and cold stress. Faeces group: formula-fed with bacterial slurry, hypoxia and cold stress. *C. butyricum* group: formula-fed with *C. butyricum*, hypoxia and cold stress. *C. tyrobutyricum* group: formula-fed with *C. tyrobutyricum*, hypoxia and cold stress.

### Histological analysis

Mouse intestinal tissue sections were soaked in 4% Carnoy’s fixative for 24h and paraffin-embedded. Then Hematoxylin-Eosin staining (Leagene, DH0006) and Alcian blue staining (Leagene, 0041) were performed on paraffin sections according to manufacturer protocol. All histological evaluation of the ileum and colon were conducted and scored in a blinded manner. The total score obtained was statistically analyzed. Alcian Blue staining was used for quantifying the coverage of mucus layer. The Ileum epithelial histological score was performed as follows [19].

Normal (score, 0); Mild (score, 1), separation of the villus core, without other abnormalities; Moderate (score, 2), villus core separation, submucosal edema, and epithelial sloughing; Severe (score, 3), denudation of epithelium with loss of villi, full thickness necrosis, or perforation.

The colon epithelial histological score was performed as follows [20].

Normal (score, 0). Excessive proliferation, abnormal crypt morphology, and goblet cell deletion (score, 1). Moderate recess loss (10-50%) (score, 2). Severe crypt loss (50-90%) (score,3). Complete absence of crypt (score,4). Small and medium-sized ulcer (ulcer surface < 10 recess length), (score, 5). Large ulcer (length of ulcer surface ≥ 10 crypts) (score,6).

### RT-qPCR analysis

Intestine tissues were homogenized using a Freeze grinder (Shanghai Jingxin) immediately after being lysed in TRIzol (Thermo Fisher, USA). For RNA was isolated by chloroform-isopropanol method and cDNA was synthesized using an iScript cDNA Synthesis Kit (Vazyme, China). cDNA was synthesized from 500 ng total RNA using HiscriptII QRT SuperMix (Vazyme, China). Real-time PCR was performed using ChamQ SYBR qPCR Master Mix (Low ROX Premixed) (Vazyme, China) and detected by ABI 7500 system (Applied Biosystems, CA, USA). Primers were used as follows:

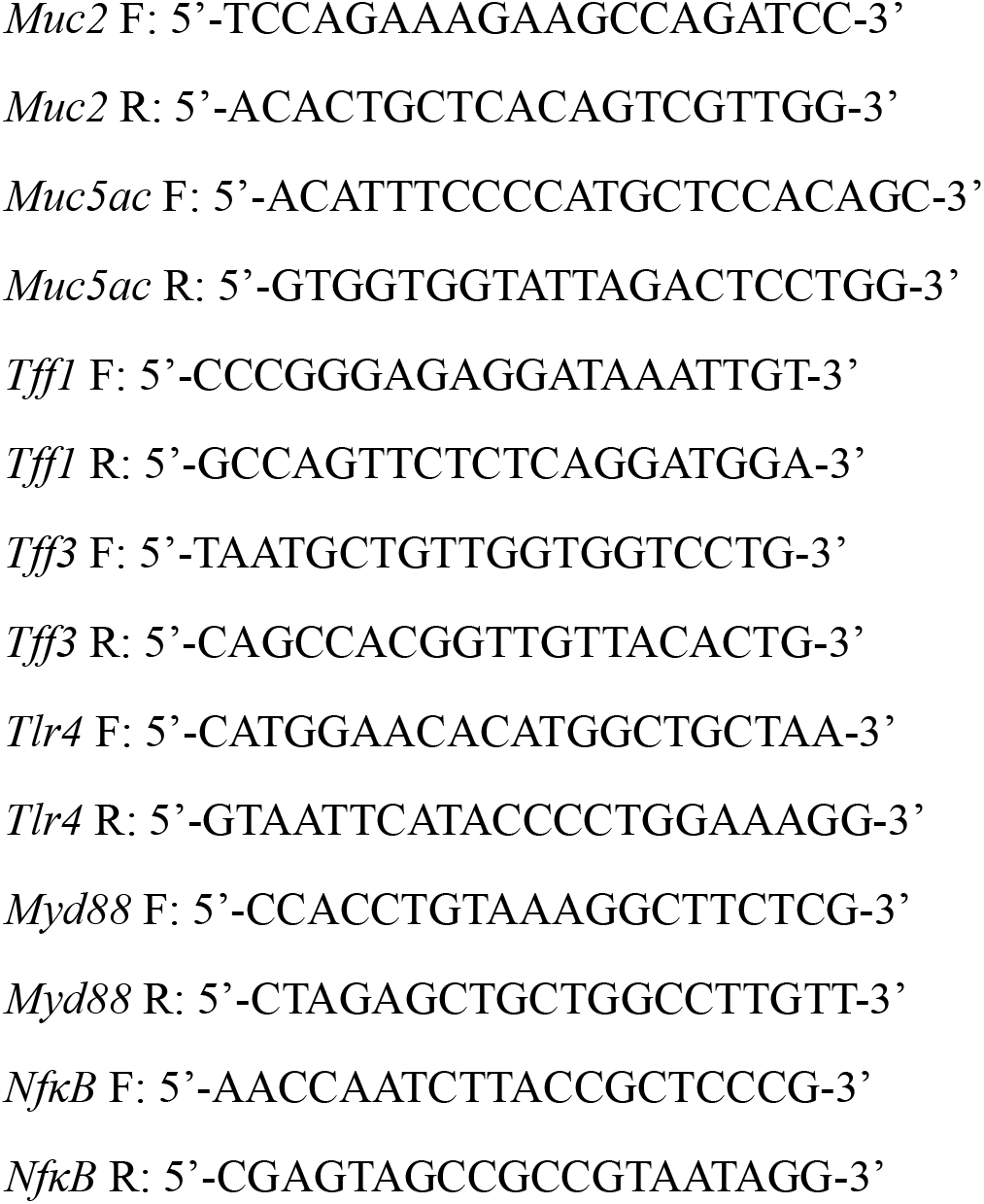

### Isolation of the lamina propria mononuclear cells (LPMCs) from ileum murine and colon

Fresh ileum and colon tissues were collected when mice were sacrificed. After being washed with cold PBS, tissues were cut into 1.5cm and incubated in 1mM dithiothreitol (DTT, Sigma-Aldrich, D9779) and 10mM HEPES solution (Sigma-Aldrich, 375368) for 10min at 37°C. After vigorous shaking and washing with PBS, 30mM ethylenediaminetetraacetic acid (EDTA, Sigma-Aldrich, 798681) and 10mM HEPES solution were used to incubate tissues twice. Then tissues were incubated in RPMI Medium 1640 (Gibco, 31800-022) with 0.2 mg/ml collagenase I (Sigma-Aldrich, C2674) and 0.15mg/ml DNase (Sigma-Aldrich, AMPD1) to digest tissues for 90 min at 37°C. After a vigorous stirrer, purified the cells by using a 70μm filter and collected them by centrifugation for 5min at 500g. A discontinuous Percoll (Cytiva, 17089102) gradient (80%/40%) was then used to separate mononuclear cells through centrifugation for 30min at 1000g without break. Collect the LPMCs at the interface between two Percoll gradients for further flow cytometry.

### Flow cytometry analysis

The cell was labeled with FITC anti-mouse CD45 Antibody (BioLegend, No.103107), APC anti-mouse/human CD11b (BioLegend, No.101211), PE/Cyanine 7 anti-mouse F4/80 (BioLegend, No. 123113), PE anti-mouse Ly-6C (BioLegend, No.128007), PerCP/Cyanine5.5 anti-mouse Ly-6G (BioLegend, No. 127616) for 20 min at room temperature. Flow cytometry analysis was performed on a CytoFLEX (Beckman coulter), and results were analyzed by FlowJo software version 10.

### Immunofluorescence imaging

Intestinal samples were fixed overnight in 4% Carnoy’s fixative. Samples were then dehydrated in gradient ethanol (100%-95%-90%-75%-50%) and Xylene solution for at least 4 h. Then 4 μm Paraffin sections were rehydrated, blocked with 3% BSA solution, and stained with antibodies overnight as follows: ZO-1 (Proteintech, 21773-AP, 1:2000), E-cadherin (Affinity, AF0131, 1:200), Claudin-1 (Affinity, DF6919, 1:200). Tissue sections were then incubated with the appropriate fluorophore-conjugated secondary antibody. Before imaging, nuclei were counterstained with DAPI or Hoechst. Fluorescence analysis was performed on a Leica SP8 Fluorescence microscope.

### Fluorescence in situ hybridization

Ileum tissues were prepared for FISH analysis by fixation in 4% Carnoy’s fixative, followed by embedding in paraffin as described above. Tissues were sectioned at a thickness of 4 µm and hybridized with a universal bacterial probe directed against the 16s rRNA gene: (EUB338 probe: 5′-GCTGCCTCCCGTAGGAGT-3’). Sample were then stained with mucin2 (Santa, sc-515032, 1:300) overnight at 4℃ and Hoechst for 10min at room temperature. Fluorescence analysis was performed on a Leica SP8 Fluorescence microscope. The percentage of bacteria was quantified per mm^2^ of ileum in each group.

### Western Bolt

Tissues were lysed in lysis buffer (P0013B, Beyotime) containing protease inhibitor cocktail (ST506, Beyotime) and phosphatase inhibitor cocktail (GB-0032, KeyGEN) in a low-temperature freeze grinder. Proteins were denatured and resolved by SDS-PAGE, then transferred onto PVDF membranes (Millipore, USA). Then the samples were probed with antibodies overnight at 4℃. After labeling with antibody and Goat Anti-Rabbit IgG (H+L) HRP antibodies (ZJ2020-R, Bioworld) and imaged by a BIORAD imaging system (chemiDOCTMXRS, Bio-Rad). Primary antibodies were used as follows: ZO-1 (Proteintech, 21773-AP, 1:5000), E-cadherin (G-10) (Santa, sc-8426,1:500), Claudin-1 (Affinity, DF6919, 1:1000), TLR4 (Santa, sc-293072, 1:500), MyD88 (CST, D80F5, 1:1000), NF-κB (Santa, sc-8008,1:500), Phospho-NF-κB (Affinity, AF2006, 1:1000), anti-GAPDH (MA5-15738, ABclonal, 1:10000).

### 16s ribosomal RNA gene sequencing

Total genomic DNA was extracted from samples using Hexadecyltrimethy Ammonium Bromide (CTAB). 16s rDNA genes were amplified using a specific primer with the barcode. All PCR reactions were carried out using Phusion Hot Start II High-Fidelity PCR Master Mix (Thermo Fisher, USA). The universal bacterial 16s rRNA gene amplicon PCR primers were used: the forward primer was 5’-CCTACGGGNGGCWGCAG-3’, and the reverse primer was 5’-GACTACHVGGGTATCTAATCC-3’. Then, the DNA monitored on 2% agarose gel was purified by means of AMPure XT beads (Beckman Coulter Genomics, Danvers, MA, USA) and quantified by Qubit (Invitrogen, USA). Sequencing libraries were generated using Agilent 2100 bioanalyzer (Agilent, USA) and Illumina (KapaBiosciences, Woburn, MA, USA). Then paired-end sequencing was performed using NovaSeq 6000 sequenator (Illumina, San Diego, CA, USA) and NovaSeq 6000 SP Reagent Kit (500 cycles) (Illumina, 20029137).

### RNA Sequencing

Total RNAs of colon were extracted using Trizol Reagent and purified by Ribo-ZeroTM Magnetic Gold Kits (Illumina, MRZG126) to remove rRNA. 3μg RNAs of each biological sample were collected to construct sequencing library using NEB Next Ultra Directional RNA LibraryPrep Kit for Illumina (NEB, Ispawich, USA). The mRNAs were measured by DEGseq software to find differential expressed genes (DEG) and DEGs were further annotated through NCBI, Uniport, GO and KEGG database.

### Cell Viability Assay

Cell viability was determined by the Cell Counting Kit-8 (CCK-8, APExBIO, K1018) assay. 1000 cells were seeded for 96-well plate in each well respectively. After being treated with different bacteria directly with multiplicity of infection of 100 for 4h under anaerobic condition, the culture medium containing bacteria was removed instead of 1640 medium with 10% FBS, 1% penicillin/streptomycin, and 100 mg/mL gentamycin for 2 hours to kill the extracellular bacteria in wells. Then the medium was replaced by 1640 medium supplemented with 10% FBS and 10% CCK-8 for further analysis.

### Bacterial competition in liquid cultures

The overnight cultures of *C. butyricum*, *C. tyrobutyricum* and *A. muciniphila* were prepared for competition experiments beforehand. For direct competition, all bacterial cultures were diluted to an initial OD_600_ of 0.1 and bacteria were centrifuged at 2000rpm for 5 min. For hot-killed bacterial experiment, *C. butyricum* and *C. tyrobutyricum* cultures were placed in an air oven at 70°C for 30min to be killed completely. Then *C. butyricum* and *C. tyrobutyricum* were inoculated with *A. muciniphila* respectively resuspended in a mixed culture medium (half RCM and half BHI with 0.1% mucin), whereas co-culture of *C. butyricum* + *A. muciniphila* and *C. tyrobutyricum* + *A. muciniphila* were tested with an initial OD_600_ of 0.05 of each strain. For bacterial indirect competition, After the *C. butyricum* and *C. tyrobutyricum* cultures grew overnight at an initial OD_600_ of 0.1in a mixed culture medium, the conditioned medium supernatant was collected by centrifuging at 4000rpm for 10 min. Then conditioned mediums were used to culture *A. muciniphila* respectively with an initial OD_600_ of 0.05 of *A. muciniphila*. Bacterial cultures were collected at 0h, 24h, 48h and 72h of each experiment simultaneously. Total bacterial DNA were extracted using Bacterial DNA Kit (OMEGA, D3350-01). qPCR on a 7500 Sequence Detector (Applied Biosystems, CA, USA) was used to calculate the number of *C. butyricum, C. tyrobutyricum* and *A. muciniphila* 16s rDNA gene copies in the co-culture medium. The primers were as followed:

*Akkermansia muciniphila* F: 5’-GTTCGGAATCACTGGGCGTA-3’

*Akkermansia muciniphila* R: 5’-CGCATTTCACTGCTACACCG-3’

### Statistical analysis

Statistical significance was determined with t-test, one-way ANOVA or two-way ANOVA test, as indicated in the figure legends. Error bars indicate SEM on all graphs. Prism 9.0 (Graphpad, La Jolla, CA) was used for all statistical analyses.

## Results

### Gut microbiota took part in the development of NEC

To clarify whether dysbiosis of gut microbiota has a profound association with the development of NEC, we compared the gut microbiota composition of NEC infants with normal neonates by utilizing 16S rDNA sequencing. Despite the alpha diversity represented no significant alteration (Figure 1A), the NMSD analysis indicated the distinct change of microbiota composition in beta diversity between NEC group and Control group (Figure 1B). The differential taxonomic similarity of intestinal bacteria at phylum level from each group reflected in *Actinobacteriota*, *Firmicutes* and *Proteobacteria* (Figure 1C). Notably, bubble plot analysis also confirmed a significant alteration in genus level (Figure 1D).

To further figure out the mechanism underlying, we used a mouse model to induce NEC, including formula feeding, hypoxia and cold stress [21] and treated mice with bacterial stock (12.5μl stool slurry in 1ml formula) cultured from stool of NEC patients for one week (Figure 1E). Considering that NEC infants are likely to present with pneumatosis cystoides intestinalis[22], we compared the severity pathology of each group and calculated the number of pneumatosis cystoides intestinalis (Figure 1F-1G). The result shown that bacterial treatment facilitated the luminal gas trapping of NEC mice. Moreover, ileum and colon tissue damage were more severe after bacteria gavage (Figure 1H) judging from H&E staining. Taken together, our data manifested the microbiome disorder promoted the development of NEC.

**Figure. 1.**
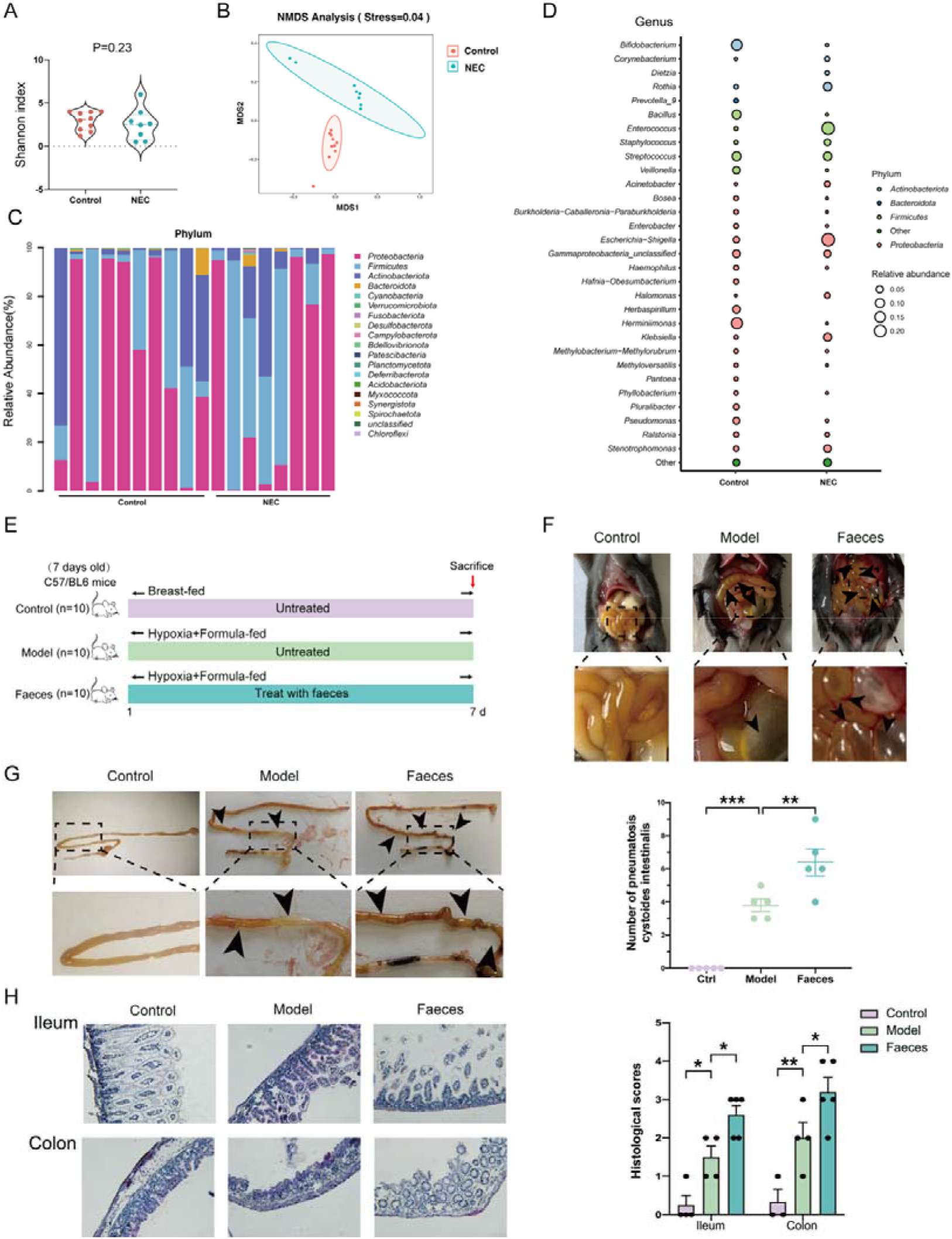
Gut microbiota took part in the development of NEC. (A) Shannon index of bacteria in human feces (n=8-10). (B) NMDS analysis of bacteria in human feces using Bray-Curtis metric distances of beta diversity. (C-D) Relative abundance of bacteria composition at phylum and genus level. (E) Experimental design for transplanting faeces from NEC patients to mice under NEC model. (F-G) Representative images of enteric cavity and canal at 7 days. The number of pneumatosis cystoides intestinalis marked by arrows were counted and shown as scatter plots (n=5). (H) H&E staining and Histological scores of ileum and colon (n=3-5). Quantified results were shown as mean ± SEM. *p*-values were generated by one-way ANOVA with multiple comparisons. *p<0.05, **p<0.01,*** p<0.001.

### *C.tyrobutyricum* protected against NEC while *C.butyricum* worsened NEC

Recent studies including epidemiological studies [23], clinical signs[24] and animal models[25] have demonstrated provenly the participation of Clostridia in NEC development. We fed NEC mice with *C.tyrobutyricum* or *C.butyricum* (both 10^9^ CFUs/ml) mixed formula as described before[21] to confirm whether these two probiotics of *clostridium* can be used for NEC treatment (Figure 2A). We found mice treated with *C.tyrobutyricum* had lighter symptom of pneumatosis cystoides intestinalis. By contrast, the symptoms of mice treated with *C.butyricum* got more severe (Figure 2B-2C). Furthermore, H&E staining results also verified that the ileum and colon damage were alleviated by *C.tyrobutyricum* but aggravated by *C.butyricum* (Figure 2D).

Briefly, two different probiotics from *clostridium* exerted reversely effects on development of NEC, which implied each bacteria strain owned unknown mechanisms behind that impacted the disease.

**Figure. 2.**
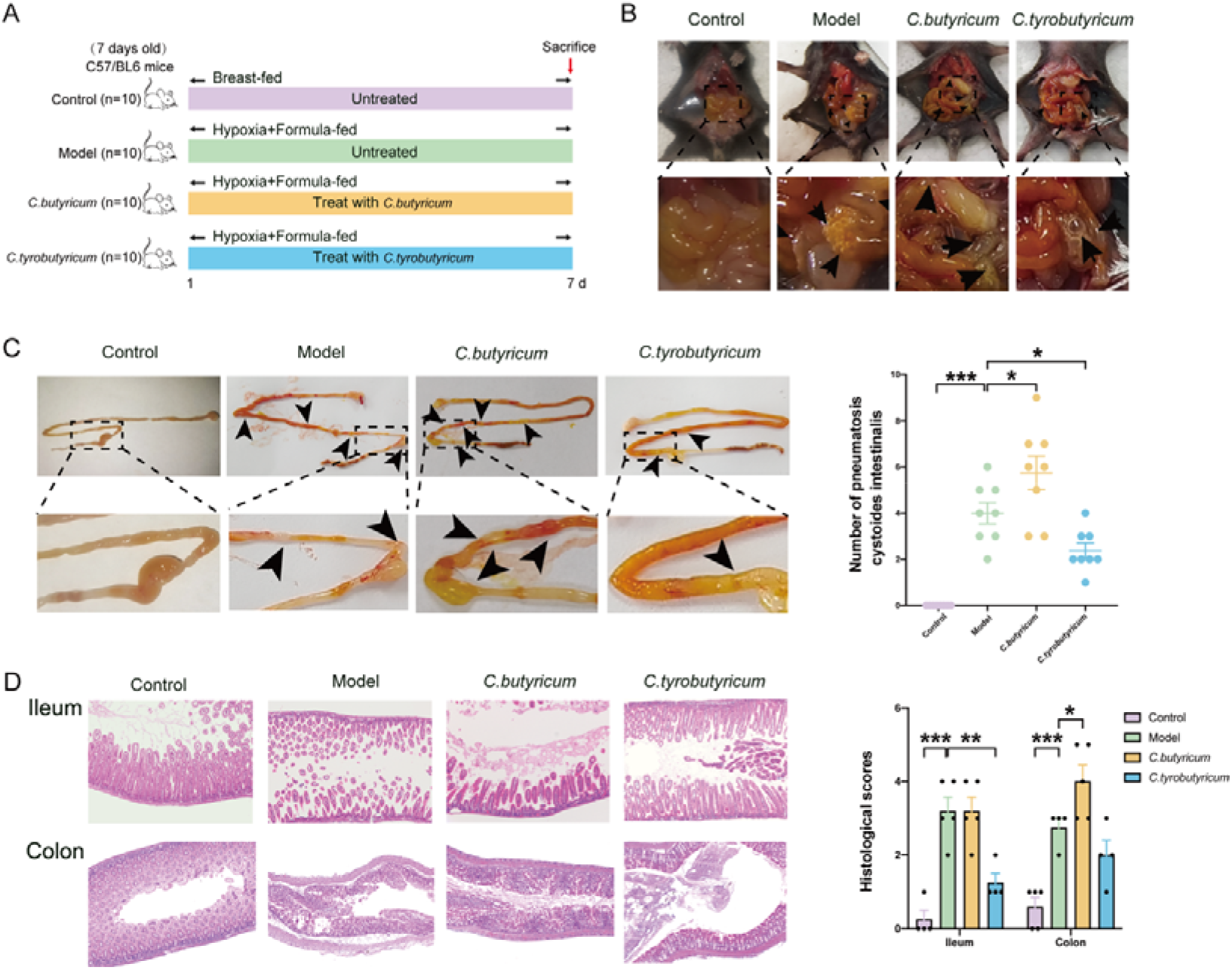
*C.tyrobutyricum* protected against NEC while *C.butyricum* worsened NEC. (A) Experimental design for treating mice with *C.butyricum* (1x10^9^ CFU/d) and *C.tyrobutyricum* (1x10^9^ CFU/d) under NEC model. (B-C) Representative images of enteric cavity and canal at 7 days. The number of pneumatosis cystoides intestinalis marked by arrows were counted and shown as scatter plots (n=8) (D) H&E staining and Histological scores of ileum and colon (n=3-5). Quantified results were shown as mean ± SEM. *p*-values were generated by one-way ANOVA or two-way ANOVA with multiple comparisons. *p<0.05, **p<0.01,*** p<0.001.

### Intestinal inflammation was alleviated by *C.tyrobutyricum* but aggravated by C.butyricum

In order to investigate the mechanism underlying both *Clostridia* species for NEC progression, we first paid attention to the alterations at transcriptional level of disease itself. In keeping with this, RNA sequencing was performed to assess messenger (m)RNA expression between NEC and normal neonate colon tissues considering that the usual sites of NEC were distal ileum and proximal colon [26]. Unsurprisingly, there was a great difference in mRNA expression comparing two groups (Figure 3A), in which 1970 upregulated mRNAs were identified and we labeled differential inflammatory-related genes in volcano plot partially (Figure 3B). What’s more, KEGG pathway enrichment emphasized that Toll-like receptor signaling pathway and NF-kappa B signaling pathway were upregulated significantly (Figure 3C), which implied an inflammation state existing in NEC neonates.

To further verify the activation of intestinal inflammation, we extracted immune cells from lamina propria layer of ileum and colon [27]. The number of CD11b^+^ F4/80^+^ macrophage, CD11b^+^ LY6c^+^ monocyte and CD11b^+^ LY6g^+^ neutrophil was detected by flow cytometry in each group mice. Notably, *C.tyrobutyricum* treatment decreased the number of macrophage, monocyte and neutrophil increased by NEC while *C.butyricum* treatment failed to invert it (Figure 3D and S1A). Consistent with RNA-sequencing data, the expression of genes in intestine tissues such as *Tlr4*, *Myd88*, *Nf-κb* and *Il-1β* were upgraded and reversed by *C.tyrobutyricum* significantly but not *C.butyricum* (Figure 3E). We examined the protein level of intestine tissues including TLR4, MyD88, NF-κB and phospho-NF-κB by western bolt (WB). Treatment with *C.butyricum* elevated the expression of TLR4, MyD88 and phospho-NF-κB, while TLR4 and phospho-NF-κB had been reduced when treated with *C.tyrobutyricum* (Figure 3F). Collectively, our data suggested that NEC mice got a severe intestinal inflammation. Two different *Clostridia* exerted exactly the opposite effects, which *C.tyrobutyricum* alleviated but *C.butyricum* aggravated the inflammation through modulating immune cells and TLR4/NF-κB signaling pathway.

**Figure. 3.**
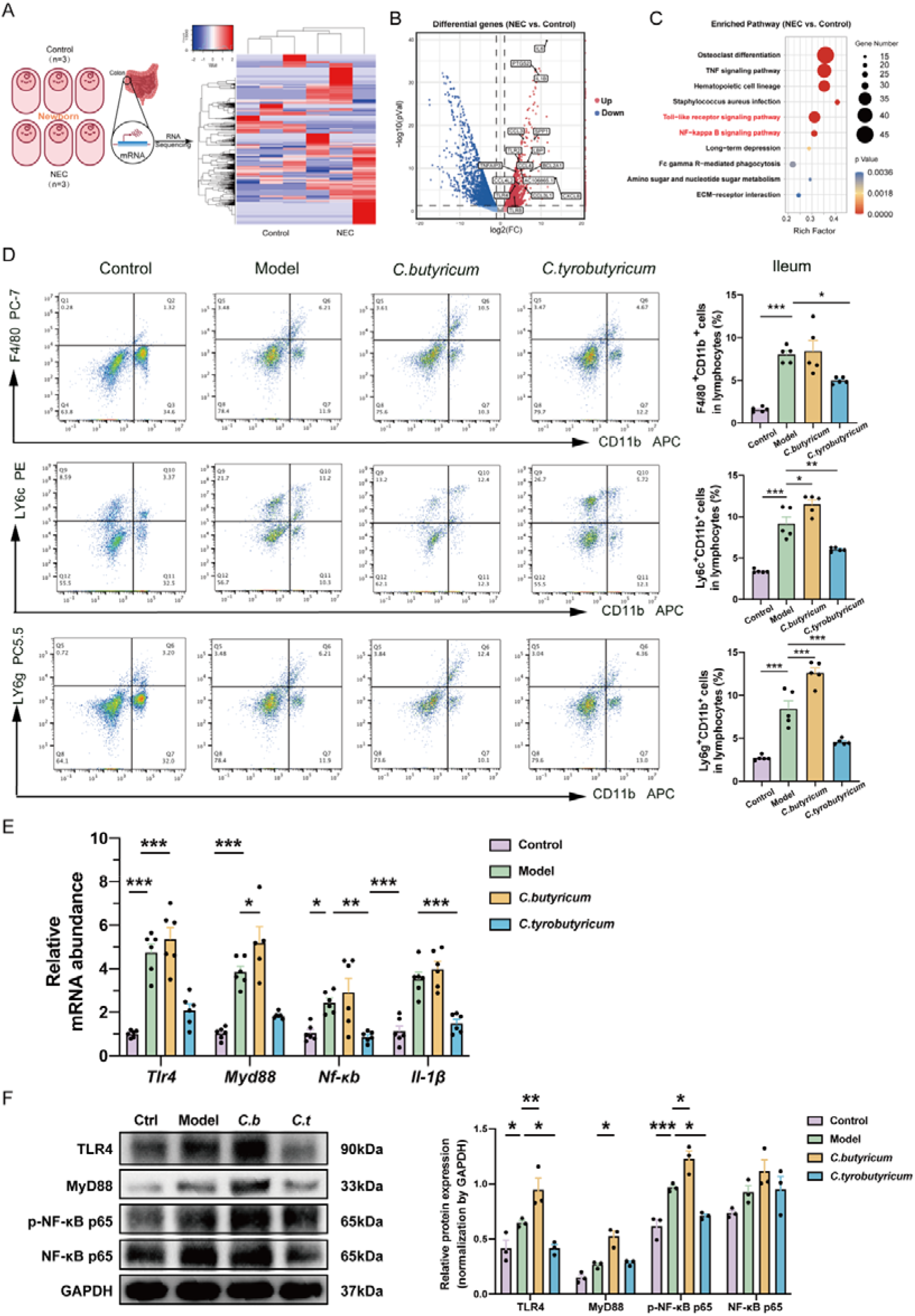
Intestinal inflammation was alleviated by *C.tyrobutyricum* but aggravated by *C.butyricum*. (A) Schematic diagram demonstrating the workflow of RNA-seq for colons of NEC. (B) Volcano plot showing the differential genes and labeling some genes upgraded in comparison between NEC mice and control mice. (C) Analysis of top 10 up-regulated pathways in NEC mice compared with control mice. (D) Flow cytometric analysis of CD11b^+^F4/80^+^(Macrophages), CD11b^+^Ly6C^+^ (Monocytes) and CD11b^+^Ly6G^+^ (Neutrophils) cells in the ileum tissues of mice (n=5). (E) mRNA expressions of *Tlr4*, *Myd88*, *Nf-κb* and *Il-1β* in mice intestine tissues with *C.butyricum* or *C.tyrobutyricum* treatment. (F) Protein level of TLR4, MyD88, phospho-NF-κB and NF-κB in intestine tissues. Quantified results were shown as mean ± SEM. p-values were generated by one-way ANOVA or two-way ANOVA with multiple comparisons. *p<0.05, **p<0.01,*** p<0.001.

### Intestinal barrier integrity was protected by *C.tyrobutyricum* but disrupted by C.butyricum

RNA-sequencing data was further analyzed to determine the alteration between NEC and normal neonates at the level of mRNA in colon. Gene set enrichment analysis (GESA) of the differentially expression genes between NEC and normal neonates detected enrichment of downregulated genes characteristic of tight junction (Figure 4A). We labeled the genes related to tight junction of 2073 downregulated genes in volcano plot (Figure 4B). Similarly, KEGG pathway enrichment emphasized that Tight junction signaling pathway was inhibited obviously, indicating the damage of intestinal barrier integrity in NEC neonates (Figure 4C).

To investigate the link between intestinal barrier integrity and the opposite effects of two *Clostridia* on the development of NEC, Alcian Blue staining was performed to observe the mucus layer thickness of ileum (Figure 4D). Statistically, treatment with *C.tyrobutyricum* improved the thickness of mucus layer while *C.butyricum* reduced it significantly. Furthermore, more bacteria were translocated into the submucous layer in ileum of NEC mice and treatment with *C.butyricum* failed to alleviate the permeability of the intestinal barrier, comparing with the dramatically protection of *C.tyrobutyricum* treatment (Figure 4E). Meanwhile, The expression of tight junction protein ZO-1 and claudin-1 and cell adhesion protein E-cadherin of ileum and colon were reduced in NEC mice through immunofluorescence and western blot, and *C.tyrobutyricum* treatment improved the expression of ZO-1 and E-cadherin remarkably (Figure 4F-4G and S2A). We also examined the mRNA level of *Muc2*, *Muc5ac*, *Tff1* and *Tff3* which were associated with mucus barrier integrity[28]in intestine tissues through RT-PCR (Figure 4H). The result further confirmed the protective effect of *C.tyrobutyricum* on mucus barrier. All in all, these data suggested that intestinal barrier integrity was protected by *C.tyrobutyricum* but disrupted by *C.butyricum*.

**Figure. 4.**
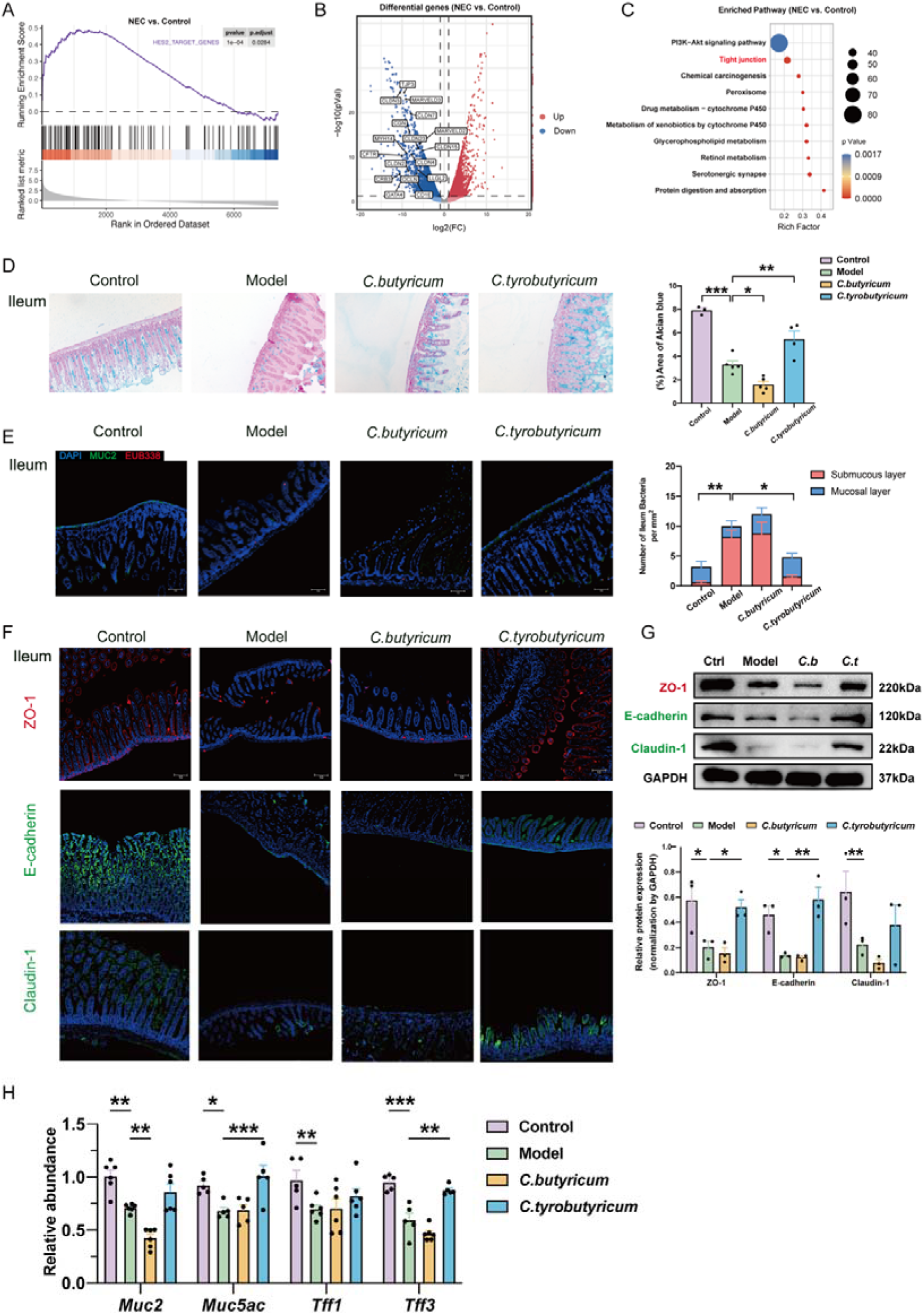
Intestinal barrier integrity was protected by *C.tyrobutyricum* but disrupted by *C.butyricum*. (A) GESA plot of genes enriched in NEC mice versus control mice was done with MSigDB (C3). (B) Volcano plot showing the differential genes and labeling some genes downgraded in comparison between NEC mice and control mice. (C) Analysis of top 10 down-regulated pathways in NEC mice compared with control mice. (D) Representative Alcian Blue images of ileum and its analysis of each group is shown as a histogram. (E) Bacteria were stained with EUB338 (red), MUC-2 (green), and DAPI (blue) by fluorescent in situ hybridization (FISH) in each group of ileum, scale bar:100μm. (F) Immunofluorescence analysis on ZO-1, E-Cadherin, and Claudin-1 in ileum sections from different groups. Representative images were shown. Scale bar: 100 µm. (G) Protein level of ZO-1, E-Cadherin, and Claudin-1 in mice intestine sections. (H) mRNA expressions of *Muc2*, *Muc5ae*, *Tff1* and *Tff3* in mice intestine tissues. Quantified results were shown as mean ± SEM. *p*-values were generated by one-way ANOVA or two-way ANOVA with multiple comparisons. *p< 0.05, **p<0.01,*** p<0.001.

### The positive effect of *C.tyrobutyricum* and the negative effect of *C.butyricum* on NEC were associated with modulating the level of *A. muciniphila*

To explore the potential association of gut microbiota dysbiosis with NEC development and examine the effects of two *Clostridia* on microbiome composition, we performed bacterial 16s rDNA sequencing in feces of mice. Although the bacteria Shannon index of four groups did not alter significantly (Figure S3A), three-dimension principal component analysis (PCA) revealed a distinct separation of microbiota for Control, Model, *C.butyricum* and *C.tyrobutyricum* group (Figure 5A). To further investigate the overall bacterial composition among each group, we compared the top 10 relative abundance of bacteria at the phylum level in different groups (Figure 5B). Consistent with previous study[11], the levels of *Bacteroidetes*, *Proteobacteria* and *Firmicutes* altered obviously. Moreover, we saw a decrease in *Verrucomicrobia* at NEC and *C.butyricum* groups, and an increase at *C.tyrobutyricum* group. The analysis of genus confirmed the decline of *Akkermansia* in NEC group and the improvement of in *Akkermansia* in *C.tyrobutyricum* group (Figure 5C and S3B). The result of linear discriminant analysis effect size (LEfse) analysis also manifested the phenomenon in mice feces (Figure 5D). Consistently, the abundance of *Akkermansia* decreased obviously in neonate samples, too (Figure 5E). Up to now, *A*. *muciniphila* is the only species of genus *Akkermansia* has ever been found. As a mucus-consuming bacterium in the intestine, the reduce of *A. muciniphila* is robust in association with multiple diseases [29]. Proper supplement of *A. muciniphila* has proven efficacy to improve intestinal barrier integrity and host immune [30]. So we measured the relative abundance of *A. muciniphila* by using qPCR in different mice groups and found the same variation trend in mice feces (Figure 5F). All in all, *C.butyricum* treatment worsened the decline of *A. muciniphila* caused by NEC model but *C.tyrobutyricum* treatment increased the abundance of *A. muciniphila* in mice feces.

**Figure. 5.**
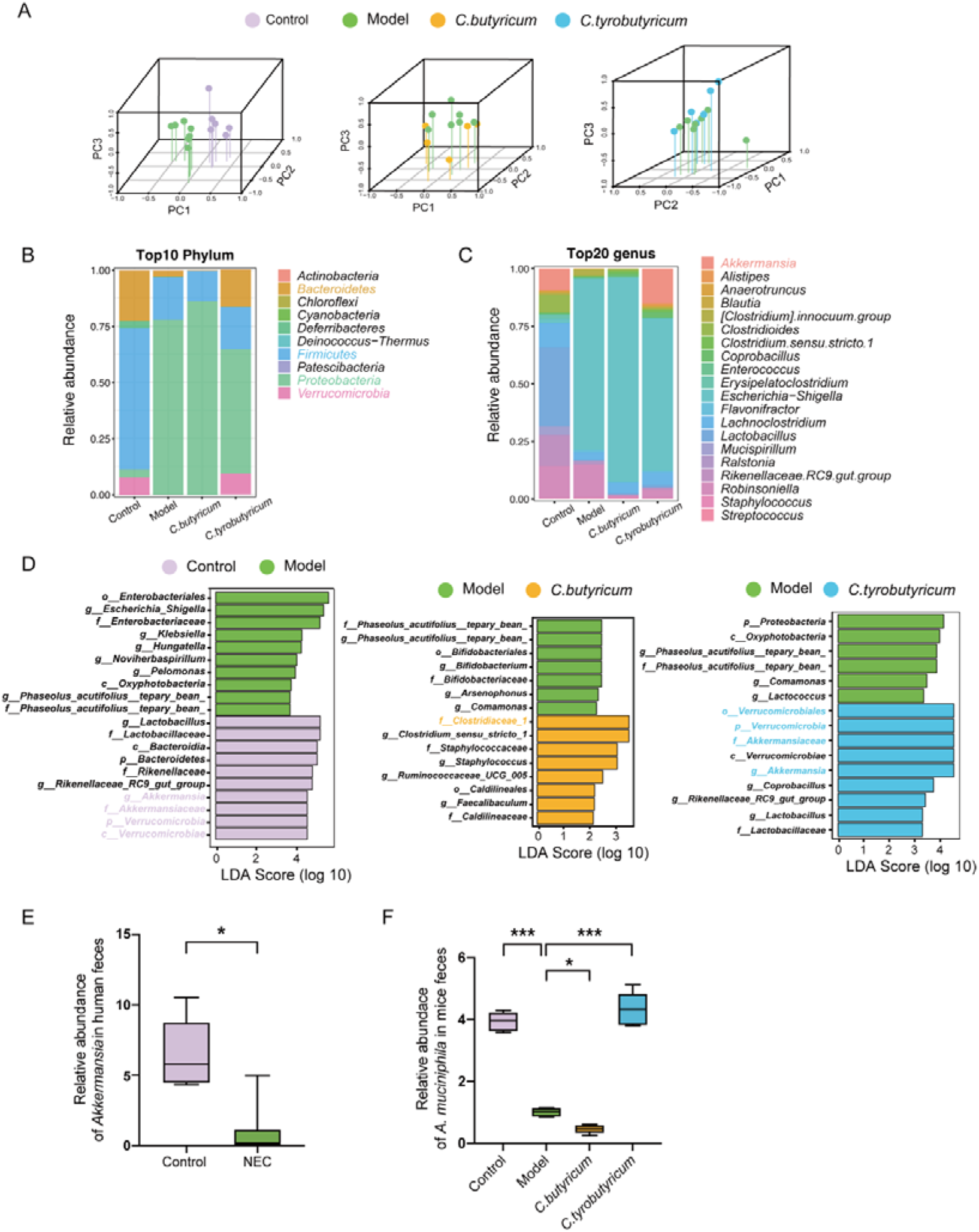
The positive effect of *C.tyrobutyricum* and the negative effect of *C.butyricum* on NEC were associated with modulating the level of *A. muciniphila*. (A) Representative images of three-dimension principal component analysis (PCA) of Control, model, *C.butyricum* and *C.tyrobutyricum* mice. (B-C) The relative abundances of bacteria at phylum and genus level were shown as a stacked bar plot. Each column corresponds to one group. (D) Representative images of linear discriminant analysis effect size (LEfSe) of each two groups. (E) Relative abundance of *Akkermansia* in neonate feces analyzed by Kruskal-Wallis test. (F) Relative 16s rDNA expression of *A. muciniphila* measured by quantitative PCR analyzed by one-way ANOVA. Quantified results were represented using box and whisker plots. p-values were generated by one-way ANOVA with multiple comparisons. ***p< 0.001.

### Interspecific competition provided a fitness advantage to *C.butyricum* over *A. muciniphila*

To further investigate the reason behind, we wondered that whether interspecific colonization resistance between the two *Clostridia* and *A. muciniphila* accounting for the variation of *A. muciniphila*. Therefore, overnight liquid cultures of each strain were prepared for bacterial competition [31]. We compared the growth rates of *A. muciniphila* directly with *C.butyricum* and *C.tyrobutyricum* under nutrient-limited conditions through absolute quantitative PCR (Figure 6A). The results depicted that *A. muciniphila* had lower growth rates when co-cultured with *C.butyricum* than proliferating singly while the yields didn’t change obviously when co-cultured with *C.tyrobutyricum*. Meanwhile, the hot-killed *C.butyricum* also suppressed the growth of *A. muciniphila* (Figure 6B), but the conditioned medium of *C.butyricum* exerted no effect on the growth of *A. muciniphila* (Figure 6C), which suggested that the colonization resistance of *C.butyricum* to *A. muciniphila* rooted in bacterial structure itself and the fermentation products or metabolites of *C.butyricum* and *C.tyrobutyricum* did not act as stimulatory molecules. Taken together, our data found that *C.butyricum* provided a more fitness advantage over *A. muciniphila* in interspecific competition.

**Fig. 6.**
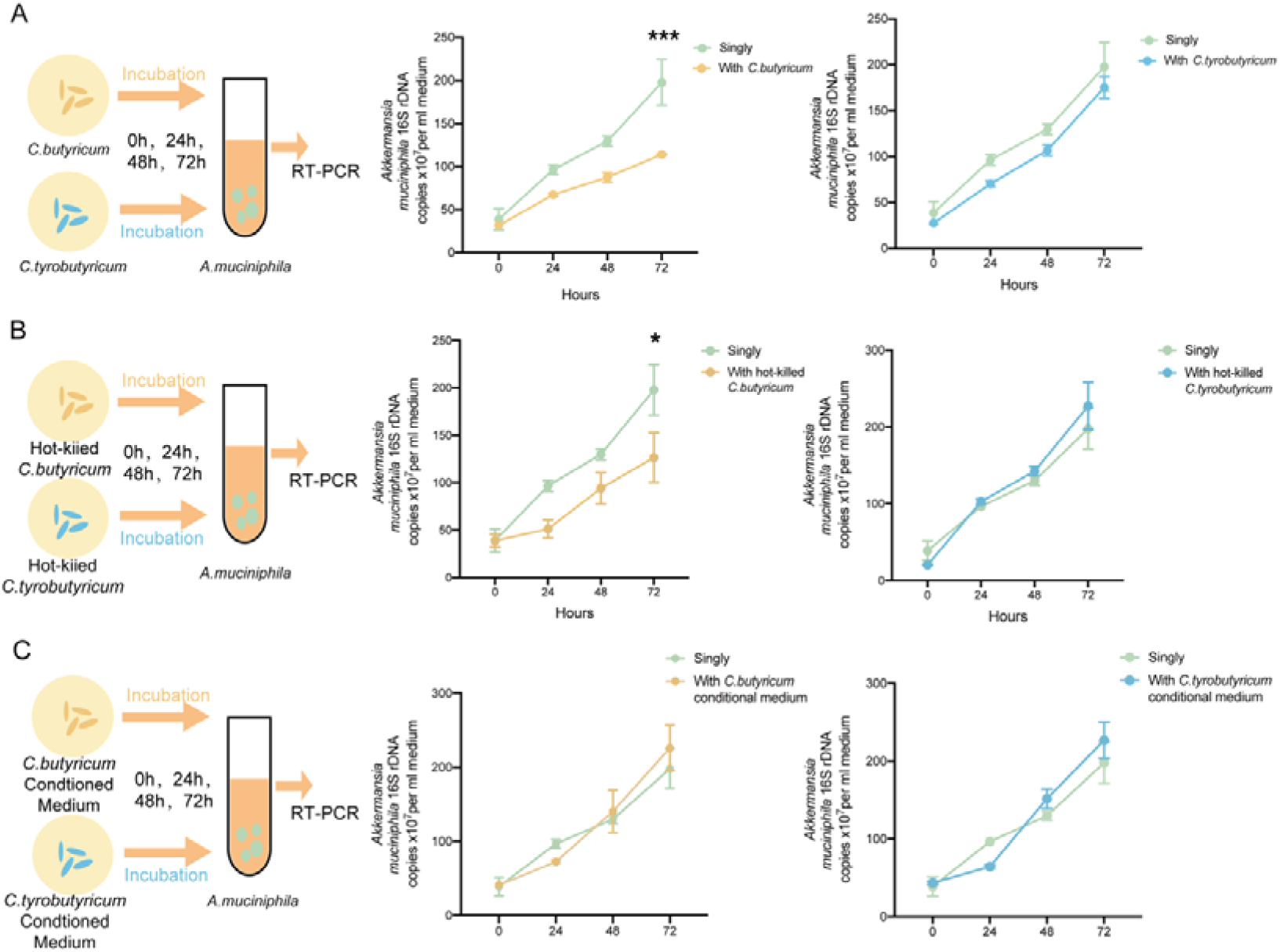
Interspecific competition provided a fitness advantage to *C.butyricum* over *A. muciniphila.* (A-C) Schematic diagram demonstrating the workflow of competition experiments in liquid cultures among different strains and the growth rates of *A. muciniphila* in each co-culture solution were represented by line charts. The growth rates of *A. muciniphila* were shown as the 16S rDNA copies measured by quantitative PCR. Quantified results were shown as mean ± SEM. *p*-values were generated by two-way ANOVA with multiple comparisons. *p<0.05, **p<0.01,*** p<0.001.

## Conclusion

NEC has become one of the health-threatening diseases in infants, especially premature neonates. For lack of comprehensive insights into pathogenesis and mechanisms underlying NEC[32], there remains an urgent need for finding out a more effective and safe therapeutic schedule. Given that gut microbiome play a key role in the healthy growth of infants including immune and intestinal barrier development and new insights also pointed out the gut microbiome dysbiosis happened before the occurrence of NEC [33]. Meanwhile, our 16S rDNA data and experimental design also confirmed the participation of gut microbiota in NEC development. Of all the therapies for intervening gut microbiome dysbiosis in stock, probiotics therapy have been proven beneficial for restoring intestinal homeostasis to reduce the incidence of NEC [34, 35]. However, do all the probiotic products apply to the NEC treatment? We found an interesting phenomenon: there was still an argument on the influence of *Clostridia* to the NEC development and *Clostridium butyricum*, a butyric acid-producing probiotic, has been associated with the NEC intimately [12].

Here, we choose two probiotics of C*lostridium*: *Clostridium butyricum* and *Clostridium tyrobutyricum* to clarify whether they are entitled to treat NEC considering the controversial issue on the effect of *Clostridia* in NEC development. NEC model was established and mice were fed with exclusive formula mixed with *C. butyricum* or *C. tyrobutyricum*. Surprisingly, the *C. tyrobutyricum* treatment alleviated the severity of NEC model while *C. butyricum* treatment aggravated the condition. Combining with our RNA-sequencing results, the NEC neonates possessed a more intense inflammatory signaling in intestines [36] and a lower intestinal barrier integrity [37]. Simultaneously, *C. tyrobutyricum* treatment could relieve the inflammation and improve the intestinal barrier integrity, but *C. butyricum* treatment worsened the condition based on our data. We further investigated the bacterial diversity and homeostasis of human and mice feces to find potential pathogenic mechanism. The 16S rDNA sequencing results indicated that the decline of *A. muciniphila* was in association with NEC development. Our finding supported that *C. tyrobutyricum* treatment could restore the level of *A. muciniphila* but *C. butyricum* treatment exacerbated the loss of *A. muciniphila*. In addition, the fitness advantage of *C. butyricum* in interspecific competition may account for the reduce of *A. muciniphila*.

Although our study revealed a clear role of *C. butyricum* and *C. tyrobutyricum* in participating NEC progression, there are still some deficiencies centralizing on this study remaining unsolved. For instance, we did not quantify the species level of *C. butyricum* and *C. tyrobutyricum* in mice feces to present the microbiota composition in vivo more detailedly. Besides, the negative result of competition between *C. tyrobutyricum* and *A. muciniphila* implied another potential mechanism worthy to be explored for the fitness advantage of *A. muciniphila* under *C. tyrobutyricum* treatment.

In conclusion, our study strongly support the idea that treatment with *C. tyrobutyricum* but not *C. butyricum* is entitled to protect against NEC by alleviating intestinal inflammation and improving intestinal barrier integrity. Meanwhile, two kinds of treatment exert opposite effects on modulating the level of *A. muciniphila*, which is deeply associated with NEC development. Therefore, *C. tyrobutyricum* supplement may have the potential to be a therapeutic strategy for NEC in clinical treatment.

**Fig. 7.**
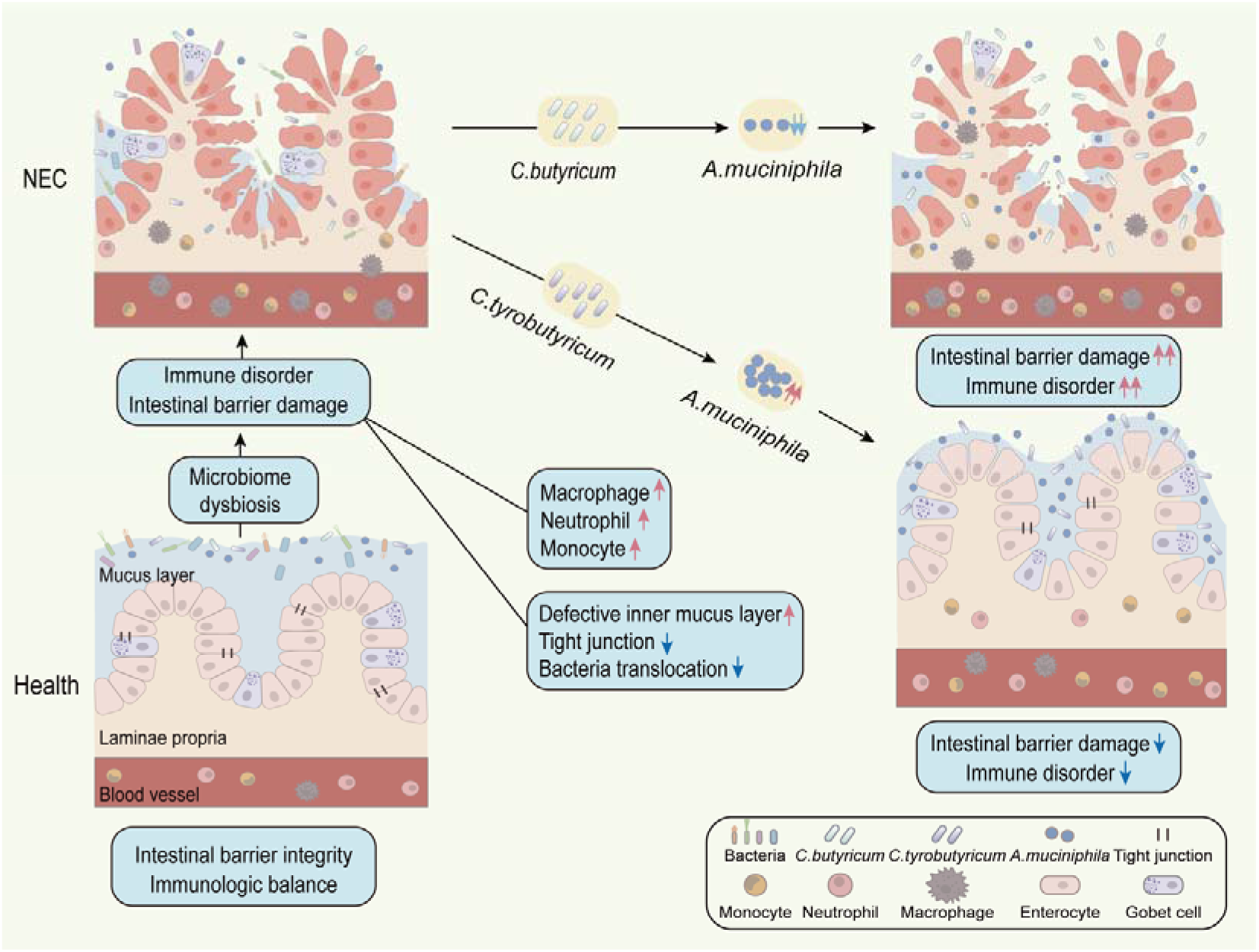
Graphical Abstract. *Clostridium butyricum* and *Clostridium tyrobutyricum* affect the development of NEC by regulating the relative abundance of *Akkermansia muciniphila*.

## Supplementary Information

**Additional file 1: Table. S1** Information of human samples related to materials and methods.

**Additional file 2: Figure. S1** Intestinal inflammation was alleviated by *C.tyrobutyricum* but aggravated by *C.butyricum*. (A) Flow cytometric analysis of CD11b^+^F4/80^+^(Macrophages), CD11b^+^Ly6C^+^ (Monocytes) and CD11b^+^Ly6G^+^ (Neutrophils) cells in the colon tissues of mice (n=4). Quantified results were shown as mean ± SEM. *p*-values were generated by one-way ANOVA with multiple comparisons. *p<0.05, **p<0.01,*** p<0.001.

**Additional file 3: Figure. S2** Intestinal barrier integrity was protected by *C.tyrobutyricum* but disrupted by *C.butyricum*. (A) Immunofluorescence analysis on ZO-1, E-Cadherin, and Claudin-1 in colon sections from different groups. Representative images were shown. Scale bar: 100 µm.

**Additional flie 4: Figure S3** The positive effect of *C.tyrobutyricum* and the negative effect of *C.butyricum* on NEC were associated with modulating the level of *A. muciniphila*. (A) Representative images of Shannon index of Control, model, *C.butyricum* and *C.tyrobutyricum* mice. (B) Heatmaps of bacteria of different groups at genus level. Each column corresponds to one sample.

## Acknowledgments

Thanks are due to Jiangning affiliated Hospital of Nanjing Medical University and Nanjing Children’s Hospital for assistance with the project. The authors thank Nanjing Jiangbei New Area Biopharmaceutical Public Service Platform for analyzing the sequencing data.We appreciate for the instruction and help from Pro. Lu and Doctor. Wei, our mentors in life and research.

## Author’ Contributions

Zhonghong Wei and Yin Lu designed the study protocol, supervised all parts of the project, and reviewed final version approval. Weibing Tang contributed to the design of the clinical trial and the analysis of clinical data. Ruizhi Tao and Gangfan Zong performed the experiments and analyzed the data. Hongxing Li and Shishan Xia were responsible for the collection and analysis of clinical samples.Yehua Pan, Peng Cheng, and Rui Deng contributed reagents/materials/analysis tools. Ruizhi Tao, Gangfan Zong, and Zhonghong Wei edited the manuscript. Wenxing Chen and Aiyun Wang checked data and provided suggestions. Yin Lu contributed to text revision and discussion. All authors read and approved the final manuscript.

## Availability of data and materials

Raw 16S rRNA sequencing data have been deposited in Sequence Read Archive (https://submit.ncbi.nlm.nih.gov/subs/sra/) with BioProject ID: PRJNA894504. The other data are available from the corresponding author upon reasonable request. All data needed to evaluate the conclusions in the paper are present in the paper. Additional data related to this paper may be requested from the authors.

## Declarations

### Ethics approval and consent to participate

All experimental protocols were approved by the Animal Care and Use Committee of the Nanjing University of Chinese Medicine (Nanjing, China) and conformed to the Guidelines for the Care and Use of Laboratory Animals (I ACUC-1908022, 05 August 2019).

### Consent for publication

Not applicable

